# The 2019 coronavirus (SARS-CoV-2) surface protein (Spike) S1 Receptor Binding Domain undergoes conformational change upon heparin binding

**DOI:** 10.1101/2020.02.29.971093

**Authors:** Courtney Mycroft-West, Dunhao Su, Stefano Elli, Yong Li, Scott Guimond, Gavin Miller, Jeremy Turnbull, Edwin Yates, Marco Guerrini, David Fernig, Marcelo Lima, Mark Skidmore

**Affiliations:** Molecular & Structural Biosciences, School of Life Sciences, Keele University, Newcastle-Under-Lyme, Staffordshire, ST5 5BG, United Kingdom; Department of Biochemistry, Institute of Integrative Biology, University of Liverpool, Crown Street, Liverpool, L69 7ZB, United Kingdom; Istituto di Ricerche Chimiche e Biochimiche G. Ronzoni, Via G. Colombo 81, 20133, Milan, Italy; School of Medicine, Keele University, Newcastle-Under-Lyme, Staffordshire, ST5 5BG, United Kingdom; School of Chemistry, Keele University, Newcastle-Under-Lyme, Staffordshire, ST5 5BG, United Kingdom

**Keywords:** heparin, coronavirus, SARS-CoV-2, COVID-19, nCoV-19, Spike, S1, RBD, circular dichroism, surface plasmon resonance, molecular modelling

## Abstract

Many pathogens take advantage of the dependence of the host on the interaction of hundreds of extracellular proteins with the glycosaminoglycans heparan sulphate to regulate homeostasis and use heparan sulphate as a means to adhere and gain access to cells. Moreover, mucosal epithelia such as that of the respiratory tract are protected by a layer of mucin polysaccharides, which are usually sulphated. Consequently, the polydisperse, natural products of heparan sulphate and the allied polysaccharide, heparin have been found to be involved and prevent infection by a range of viruses including S-associated coronavirus strain HSR1. Here we use surface plasmon resonance and circular dichroism to measure the interaction between the SARS-CoV-2 Spike S1 protein receptor binding domain (SARS-CoV-2 S1 RBD) and heparin. The data demonstrate an interaction between the recombinant surface receptor binding domain and the polysaccharide. This has implications for the rapid development of a first-line therapeutic by repurposing heparin and for next-generation, tailor-made, GAG-based antivirals.

## Introduction

Heparin, the second most widely used drug by weight globally, is formulated as a polydisperse, heterogenous natural product. Unfractionated heparin, low molecular weight heparins and heparinoids are clinically approved as anticoagulants / thrombotic with excellent safety, stability, bioavailability and pharmacokinetic profiles. Crucially, heparin and its derivatives, some of which lacking significant anticoagulant activity^1^, are an under-exploited antiviral drug class, despite possessing broad-spectrum activity against a multitude of distinct viruses, including *coronaviridae* and SARS-associated coronavirus strain HSR1^2^, in addition to flaviviruses^3,4^, herpes^5^, influenza^6^ and HIV^7,8^.

Traditional drug development processes are slow and ineffective against emerging public health threats such as the current SARS-CoV-2 coronavirus outbreak which makes the repurposing of existing drugs a timely and attractive alternative. Heparin, a well-tolerated anticoagulant pharmaceutical, has been used safely in medicine for over 80 years and alongside its anticoagulant activities, its ability to prevent viral infection, including *coronaviridae*, has been described^1^. Furthermore, the closely related glycosaminoglycan (GAG) member, heparan sulphate (HS), is known to bind CoV surface proteins and to be used by coronavirus for its attachment to target cells^9^.

Currently, there are no commercially available medicinal products designed to treat and/or prevent infections associated with the new SARS-CoV-2 coronavirus outbreak. Here, we describe preliminary tests for the ability of the SARS-CoV-2 S1 RBD to bind heparin, an important prerequisite for the underpinning research related to the development of SARS-CoV-2 heparin-based therapeutic.

## Methods & Materials

### 2.1 Recombinant expression of SARS-CoV-2 S1 RBD

Residues 330−583 of the SARS-CoV-2 Spike Protein (GenBank: MN908947) were cloned upstream of a N-terminal 6 x HisTag in the pRSETA expression vector and transformed into SHuffle® T7 Express Competent *E. coli* (NEB, UK). Protein expression was carried out in MagicMedia™ *E. coli* Expression Media (Invitrogen, UK) at 30°C for 24 hrs, 250 rpm. The bacterial pellet was suspended in 5 mL lysis buffer (BugBuster Protein Extraction Reagent, Merck Millipore, UK; containing DNAse) and incubated at room temperature for 30 mins. Protein was purified from inclusion bodies using IMAC chromatography under denaturing conditions. On-column protein refolding was performed by applying a gradient with decreasing concentrations of the denaturing agent (6 - 0 M Urea). After extensive washing, protein was eluted using 20 mM NaH2PO4, pH 8.0, 300 mM NaCl, 500 mM imidazole. Fractions were pooled and buffer-exchanged to phosphate-buffered saline (PBS; 140 mM NaCl, 5 mM NaH2PO4, 5 mM Na2HPO4, pH 7.4; Lonza, UK) using a Sephadex G-25 column (GE Healthcare, UK). Recombinant protein was stored at −20°C until required.

### 2.2 Secondary structure determination of SARS-CoV-2 S1 RBD by circular dichroism spectroscopy

The circular dichroism (CD) spectrum of the SARS-CoV-2 S1 RBD in PBS was recorded using a J-1500 Jasco CD spectrometer (Jasco, UK), Spectral Manager II software (JASCO, UK) and a 0.2 mm pathlength, quartz cuvette (Hellma, USA) scanning at 100 nm.min^−1^ with 1 nm resolution throughout the range 190 - 260 nm. All spectra obtained were the mean of five independent scans, following instrument calibration with camphorsulphonic acid. SARS-CoV-2 S1 RBD was buffer-exchanged (prior to spectral analysis) using a 5 kDa Vivaspin centrifugal filter (Sartorius, Germany) at 12,000 g, thrice and CD spectra were collected using 21 μl of a 0.6 mg.ml^−1^ solution in PBS, pH 7.4. Spectra of heparin (porcine mucosal heparin), its derivative and oligosaccharides were collected in the same buffer at approximately comparable concentrations, since these are disperse materials. Collected data were analysed with Spectral Manager II software prior to processing with GraphPad Prism 7, using second order polynomial smoothing through 21 neighbours. Secondary structural prediction was calculated using the BeStSel analysis server^10^. To ensure that the CD spectral change of SARS-CoV-2 S1 RBD in the presence of porcine mucosal heparin did not arise from the addition of the heparin alone, which is known to possess a CD spectrum at high concentrations^11,12^ a difference spectrum was analysed. The theoretical, CD spectrum that resulted from the arithmetic addition of the CD spectrum of the SARS-CoV-2 S1 RBD and that of the heparin differed from the observed experimental CD spectrum of SARS-CoV-2 S1 RBD mixed with heparin. This demonstrates that the change in the CD spectrum arose from a conformational change following binding to porcine mucosal heparin.

### 2.3 Surface Plasmon Resonance determination of SARS-CoV-2 S1 RBD binding to unfractionated heparin

Human FGF2 was produced as described by Duchesne *et al.^13^*. Porcine mucosal heparin was biotinylated at the reducing end using hydroxylamine biotin (ThermoFisher, UK) as described by Thakar *et al*. ^14^. Heparin (20 μl of 50 mg mL^−1^) was reacted with 20 μl hydroxylamine-biotin in 40 μl 300 mM aniline (Sigma-Aldrich, UK) and 40 μl 200 mM acetate pH 6 for 48 h at 37 °C. Free biotin was removed by gel-filtration chromatography on Sephadex G25 (GE LifeSciences, UK).

A P4SPR, multi-channel Surface Plasmon Resonance (SPR) instrument (Affinté Instruments; Montréal, Canada) was employed with a gold sensor chip that was plasma cleaned prior to derivatization. A self-assembled monolayer of mPEG thiol and biotin mPEG was formed by incubating the chip in a 1 mM solution of these reagents at a 99:1 molar ratio in ethanol for 24 hrs^15^. The chip was rinsed with ethanol and placed in the instrument. PBS (1X) was used as the running buffer for the three sensing and a fourth background channel at 500 μl.min^−1^, using an Ismatec pump. Twenty micrograms of streptavidin (Sigma, UK; 1 ml in PBS) were injected over the four sensor channels. Subsequently, biotin-heparin (1 ml) was injected over the three sensing channels.

Binding experiments used PBS with 0.02% Tween 20 (v/v) as the running buffer. The ligand was injected over the three sensing channels, diluted to the concentration indicated (see figures) at 500 μl.min^−1^. Sensor surfaces with bound FGF2 were regenerated by a wash with 2 M NaCl (Fisher Scientific, UK). However, this was found to be ineffectual for SARS-CoV-2 S1 RBD. Partial regeneration of the surface was achieved with 20 mM HCl (VWR, UK) and only 0.25 % (w/v) SDS (VWR, UK) was found to remove the bound protein. After regeneration with 0.25 % (w/v/) SDS, fluidics and surfaces were washed with 20 mM HCl to ensure all traces of the detergent were removed. Background binding to the underlying streptavidin bound to the mPEG/biotin mPEG self-assembled monolayer was determined by injection over the control channel. Responses are reported as the change in plasmon resonance wavelength, in nm and for the three control channels represent their average response.

## Results

### 3.1 Surface Plasmon Resonance binding studies

FGF2, a well characterised heparin-binding protein was used to test the successful functionalization of the three sensing channels with biotin-heparin. Injection of 1 mL 100 nM FGF2 over the sensing channels elicited a significant response (Fig. 1A, injection between the red arrows). However, 100 nM FGF2 elicited no response in the control channel, functionalized solely with streptavidin (not shown). The bound FGF2 was removed by a wash with 2 M NaCl, as done previously for the IASys optical biosensor^16^.

**Figure 1.**
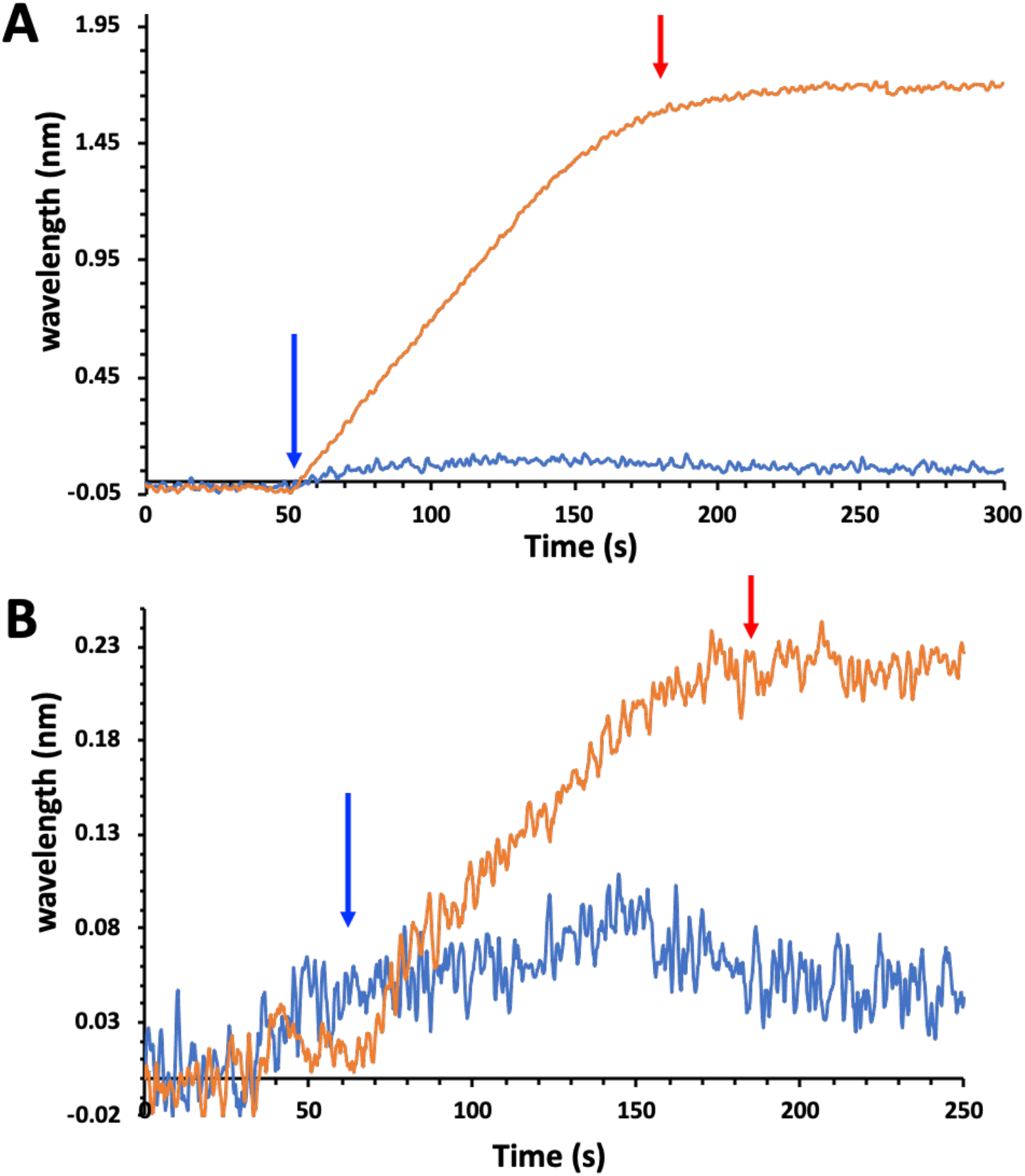
Interaction of FGF2 and 100 nM SARS-CoV-2 S1 RBD with immobilised heparin. Reducing end biotinylated heparin (-) was immobilised on a streptavidin functionalised P4SPR sensor surface (no biotin-heparin (-) control). PBS running buffer flow rate was 500 μl.min^−1^. The data for the three sensing channels are reported as an average response (-). The start of protein injections are indicated by blue arrows and the return of the surface to running buffer (PBST) by red arrows. (A) Injection of 100 nM FGF2. (B) Injection of 100 nM SARS-CoV-2 S1 RBD protein.

When 65 nM SARS-CoV-2 S1 RBD was injected over the three sensing channels, there was an initial decrease in signal, followed by an increase, indicative of binding (Fig. 1B between red arrows). The initial decrease was due to a slightly lower refractive index of the SARS-CoV-2 S1 RBD protein solution compared to the PBS of the running buffer, which caused a negative bulk shift. This is demonstrated by the injection of 65 nM solution over the control channel, functionalized with just streptavidin, where there was a decrease in response, followed by a return to baseline when the channel was returned to running buffer (Fig. 1C, between red arrows). These data demonstrate that the SARS-CoV-2 S1 RBD protein binds specifically to heparin immobilised through its reducing-end and fails to bind to the underlying streptavidin / ethyleneglycol surface.

### 3.2 Secondary structure determination of SARS-CoV-2 S1 RBD protein by circular dichroism spectroscopy

Circular dichroism (CD) spectroscopy detects changes in protein secondary structure that occur in solution using UV radiation. Upon binding, conformational changes are detected and quantified using spectral deconvolution^17^. Indeed, SARS-CoV-2 S1 RBD underwent conformational change in the presence of heparin (Figure 2). Combined, helix content increased by 1.5% and global beta-sheet content decreased by 2.1%. The observed changes further demonstrate that the SARS-CoV-2 S1 RBD interacts with heparin in aqueous solution of physiological significance, whereby the major changes induced by heparin are those associated with antiparallel and helix content.

**Figure 2.**
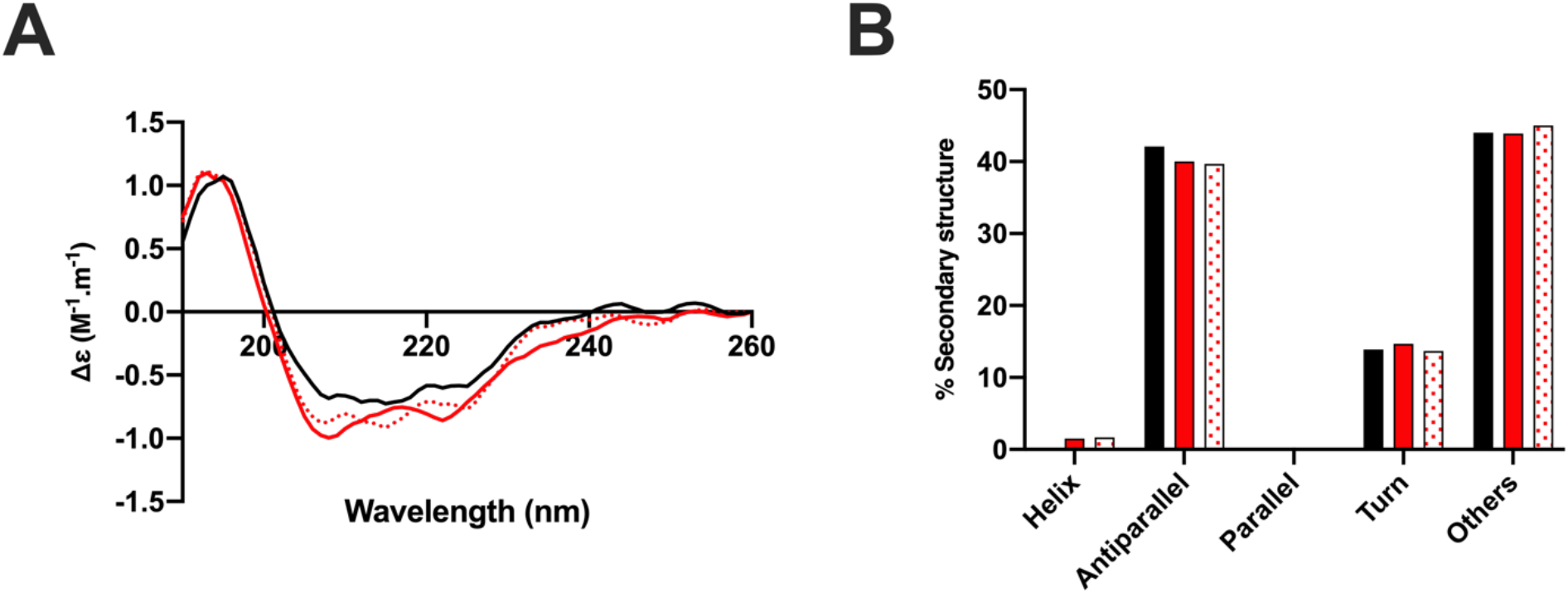
The structural change of the SARS-CoV-2 S1 RBD observed in the presence of heparin by circular dichroism (CD) spectroscopy. (**A**) CD spectra of spike 1 RBD alone (black) or with heparin (red) in phosphate buffered saline pH 7.4. Theoretical sum of spike 1 RBD alone and heparin (control) if no interaction was observed (dotted red). (**B**) Δ secondary structure (%) of (A). A 1.5% increase in helix and 2.1% decrease in antiparallel secondary structural features were calculated (BestSel) for the observed spectrum compared to that of the theoretical, summative spectrum of the SARS-CoV-2 S1 RBD in the presence of heparin.

Basic amino acids are known to dictate the binding between proteins and heparin. With that in mind, primary sequence analysis of the expressed protein domain and analysis of the modelled SARS-CoV-2 S1 RBD structure (Figure 3) shows that there are several potential heparin binding sites and, more importantly, that thesespatches of basic amino acids are exposed on the protein surface.

**Figure 3.**
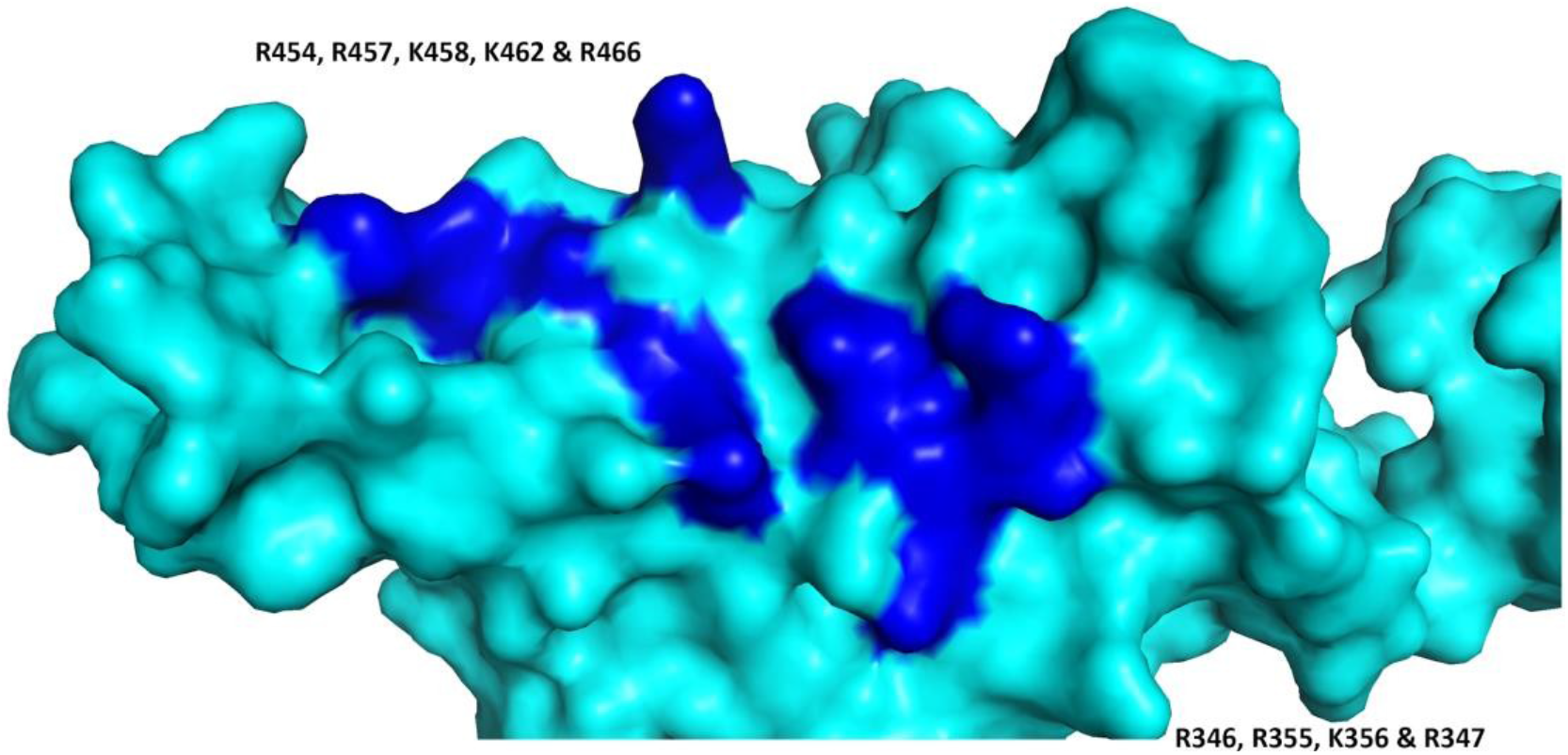
SARS-CoV-2 S1 RBD protein model. Basic amino acids that are solvent accessible on the surface are indicated (dark blue); these can be observed to form a continuous patch.

## Discussion and Conclusion

The rapid spread of SARS-CoV-2 represents a significant challenge to global health authorities and, as there are no currently approved drugs to treat, prevent and/or mitigate its effects, repurposing existing drugs is both a timely and appealing strategy. Heparin, a well-tolerated anticoagulant drug, has been used successfully for over 80 years with limited and manageable side effects. Furthermore, heparin belongs to a unique class of pharmaceuticals that has effective antidotes available, which makes its use safer.

Studying SARS-CoV-2 Spike protein structure and behaviour in solution is a vital step for the development of effective therapeutics against SARS-CoV-2. Here, the ability of the SARS-CoV-2 S1 RBD to bind pharmaceutical heparin has been studied using spectroscopic techniques in concert with molecular modelling. The data show that SARS-CoV-2 S1 RBD binds to heparin and that upon binding, a significant structural change is induced. Moreover, moieties of basic amino acid residues, known to constitute heparin binding domains, are solvent accessible on the SARS-CoV-2 S1 RBD surface and form a continuous patch that is suitable for heparin binding.

Glycosaminoglycans are ubiquitously present on almost all mammalian cells and this class of carbohydrates are central to the strategy employed by *coronaviridae* to attach to host cells. Heparin has previously been shown to inhibit SARS-associated coronavirus strain HSR1 cell invasion ^2,18^ and this, in concert with the data presented within this study, supports the utilisation of glycosaminoglycan-derived pharmaceuticals against SARS-associated coronavirus. Furthermore, this study strongly supports the repurposing of heparin and its derivatives as antiviral agents, providing a rapid countermeasure against the current SARS-CoV-2 outbreak.

It is noteworthy that even pharmaceutical-grade heparin preparations remain a polydisperse mixture of natural products, containing both anticoagulant and non-anticoagulant saccharide structures. The latter may prove to be an invaluable resource for next-generation, biologically active, antiviral agents that display negligible anticoagulant potential, whilst the former remains tractable to facile, chemical (and enzymatic) engineering strategies to ablate their anticoagulation activities.

The subfractionation of existing heparin preparations against anticoagulant activities (with proven low-toxicity profiles, good bioavailability and industrial-scale manufacturing) for off-label pathologies, provides an attractive strategy for quickly and effectively responding to COVID-19 and for the development of next generation heparin-based therapeutics.

Such drugs will be amenable to routine parenteral administration through currently established routes and additionally, direct to the respiratory tract via nasal administration, using nebulised heparin, which would be unlikely to gain significant access to the circulation. Thus, the anticoagulant activity of heparin, which can in any event be engineered out, would not pose a problem. Importantly, such a route of administration would not only be suitable for prophylaxis, but also for patients under mechanical ventilation ^19^.

